# Antiviral action of different molecules obtained from invertebrates against coronavirus

**DOI:** 10.1101/2024.01.04.574161

**Authors:** RZ Mendonça, RM Nascimento, ACO Fernandes, Maria Isabel Pinheiro de Almeida, PI da Silva Junior

## Abstract

The few availability of antivirals for new strains of highly pathogenic viruses has become a serious public health problem that leads to the death of thousands of people annually. For this reason, the search for new products against these agents has become an urgent need. Many studies have been carried out with this aim. Among the multiple sources of research for new antibiotics and antivirals, bioprospecting for molecules obtained from invertebrates, or their products, has become an increasingly frequent option. Arthropods appeared on the planet around 350 million years ago and have been one of the beings with the greatest adaptability and resistance. Invertebrates have been found in all known ecosystems. Their survival for so long, in such different environments, is an indication that they have a very efficient protection system against environmental infections, despite not having a developed immune system like mammals. Historically, products obtained mainly from bees, such as honey and propolis, have been shown to be of great pharmacologica l importance, being used as antimicrobials, anti-inflammatory, antitumor, healing and several other functions. However, these molecules have also been obtained from other invertebra tes, such as caterpillars and spiders. Previous studies by our group have demonstrated an intense antiviral and antimicrobial activity of these materials. In this study, we identified, isolated and characterized compounds with potent antiviral effect against avian Coronavirus in propolis from *Scaptotrigona aff postica*, hemolymph of *Lonomia obliqua* and from mygalin, an acylpolyamine, isolated from hemocytes of spider *Acanthoscurria gomesiana*. The antiviral assay was carried out by reducing infectious foci in cultures of infected cells and treated with these differents substances obtained from invertebrates. Propolis and crude hemolymph reduced avian coronavirus by an average of 256 x when used at a concentration of 5% v/v and an average reduction of 8x when 160µM of Mygalin was used. Propolis purified and sinthetic hemolymph reduced the virus titer by an average 64 fold. The virus reduction with synthetic mygalin, at a concentration of 26 uM, was average of 16 times. The antiviral responses of the 3 substances were dose dependent. By the other hand, the virus titer reduction was 2 times more intense when the substances were added 1 hour before cell infection with the virus. The chemical characterization of the elements present in the extracts was carried out by liquid chromatography.

## 1. INTRODUCTION

### 1.1 Coronavirus

Recently, coronavirus has become public knowledge because of the COVID 9 pandemic. This virus is not new, as we have been living with it for a long time. Coronaviruses belong to a large viral family, known since the mid-1960s, and cause mild to moderate respiratory infections in humans and animals. Generally, the clinical feature of these infections is similar to a common cold. Most people become infected with coronavir uses throughout their lives, with young children being more likely to become infected. A good percentage of alleged flu cases are actually not caused by the Influenza virus but by coronavirus. Symptoms may include runny nose, cough, sore throat and fever. These viruses can sometimes cause lower respiratory tract infections, such as pneumonia. This condition is more common in people with cardiopulmonary diseases, compromised immune systems or the elderly. However, some coronaviruses can cause severe respiratory syndromes, such as severe acute respiratory syndrome, known by the acronym SARS (Severe Acute Respiratory Syndrome). The first cases of SARS were reported in China in 2002 (Cherry, 2004). SARS-CoV quickly spread to more than a dozen countries in North America, South America, Europe and Asia, infecting more than 8,000 people and causing about 800 deaths, before of the epidemic being controlled in 2003 (MMWR, 2003). Since 2004, no cases of SARS have been reported worldwide. In 2012, another new coronavirus was isolated, distinct from the one that caused SARS at the beginning of the decade. This new coronavirus was unknown as an agent of human disease until its identification, initially in Saudi Arabia and, later, in other countries in the Middle East, Europe and Africa (Lou and Gao, 2020).

Most coronaviruses generally infect only one animal species or at least a small number of closely related species. However, periodically, a virus with common circulat ion in an animal may adapt to another species, such as humans, as is the current case with COVID-19, or a common low-risk viral strain may undergo some genetic change that makes it highly pathogenic. Coronaviruses cause epidemics not only in humans, but also in animals of commercial interest such as birds, generating high losses. These birds are affected by a coronavirus belonging to the gammacoronavirus genus, IBV - Infectious bronchitis virus. IBV and other viruses of the gamma genus do not infect humans (Yuan et al, 2021). IBV is a highly infectious disease that affects the respiratory and genitourinary tract, whether in laying or meat birds, at all ages and regions of the world. IBV spreads quickly among birds, via respiratory aerosols, feces, within a single flock or between different flocks. Transmis s ion can occur when there is contact between a sick bird and a healthy one or due to biosecurit y failures on the farm. The disease is predominantly respiratory, with cough, dyspnea, respiratory failure and suffocation (Cavanagh, 2007). Infection in the first days of life can cause mortality in young birds and, if death does not occur, symptoms tend to disappear within 10 to 15 days. In older birds, mortality is not common and, when it occurs, it is often related to co-infection with another pathogen, usually the bacterium Escherichia Coli. The IBV control in chickens is of great commercial interest. By the other hand, IBV and other viruses of the gamma genus do not infect humans. Therefore as they have a very similar replication mechanism, which can make them a good tool for studying the replication and antiviral action of substances against the human coronavirus.

### 1.2 Propolis

The composition of the word propolis comes from the Greek, pro means “in defense”, and polis means “city”, that is, “in defense of the city”. Propolis is a resinous material that bees collect from plants, mix with wax and use to build the hive. Bees apply propolis in thin layers to internal walls to strengthen and block holes and crevices, as well as using it as an “embalming” substance on dead invaders or even dead bees, thus inhibiting the growth of fungi and bacteria within the hive (Bankova et al., 2000; Ghisalberti, 1979). The medicinal properties of propolis have been reported since ancient times. In Egypt it was used in the mummification process to prevent bodies from rotting. The Greeks and Romans used it as an antiseptic and healing agent, and the Incas used it as an antipyretic (Ghisalberti, 1979, Sforcin & Bankova, 2011)

Due to its great pharmacological potential, propolis has become the target of several studies, arousing the interest of scientific society that has sought to investigate its compounds and biological activities. This interest has intensified in recent decades (Bankova et al., 2000; Araújo et al., 2010). Countries such as Switzerland and Germany already legally recognize propolis as a medicine (Bogdanov, 2014). In Brazil, the main substances described are those derived from prenylated p-coumaric acid and acetophenone, such as diterpenes and ligna ns (Bogdanov, 2014).

Studies with propolis have already demonstrated biological activities, such as immunomodulation, antitumor, antioxidant, antibacterial, antiviral and analgesic (Búfalo et al., 2009; Dobrowolski et al., 1991; Paulino et al., 2003). However, the vast majority of studies are carried out with bee propolis of commercial interest, mainly *Apis mellifera*. Studies show that epidemics in hives greatly reduce productivity (Evans and Pettis, 2005), as well as the number of bees (Moret and Schmidt, 2000). Therefore, the chemical compounds present in these products are fundamental for the protection and survival of bees, being considered an evolutionary mechanism for the benefit of the colony (Simone-Finstrom, et al., 2017). Some of the most important compounds present in resins are: **Flavonoids:** main constituents of propolis, they contribute to many biological activities (antiviral, antimicrob ia l and anti-inflammatory). Flavonoids can be classified into flavones, flavonols, flavano nes, flavanonols, chalcones, dihydrochalcones, isoflavones, isodihydroflavones, isoflavans and neoflavonoids. Brazilian red propolis contains flavonoids derived from the resin of leguminous plants (Dalbergia ecastophyllum) (Huang, et al., 2014; Zabaiou, et al., 2017); **Terpenoids:** are responsible for the characteristic resin odor in propolis and have antioxida nt and antimicrobial activities (Huang, et al., 2014; Zabaiou, et al., 2017). **Phenols:** are composed of different acids, such as cinnamic acid; demonstrate antimicrobial activit ies. Brazilian green propolis is rich in phenols (Huang, et al., 2014; Zabaiou, et al., 2017).

The composition of propolis is extremely variable, both in relation to geographic location and season (Negri et al, 2022). In Europe, North America and temperate zones, flavonoid compounds, phenolic acids and esters predominate in propolis (Falcão et al., 2010). In the Mediterranean, terpenoids have been identified in high concentrations (Popova, et al., 2010; Popova, et al., 2011). In Africa, the main compounds are triterpenoids (Zabaiou, et al., 2017). The chemical composition of propolis is influenced by both the environme ntal vegetation, the seasons and the bees that produce it (Bankova et al., 2000; Negri et al, 2022). These variations in chemical components affect their biological properties (Nakamura et al., 2010). This is why it is so important to study propolis from different species of bees and from different regions.

Among the different species of bees, there are few studies that deal with propolis from stingless bees. Only recently have these bees attracted attention and studies have been carried out to characterize their activities.

In Brazil, one of the species of stingless bees belongs to the Apidae family, from the Meliponinae subfamily. These bees mix plant resin with wax and clay, thus forming geopropolis. This name is given precisely because of the use of land for the final compositio n of the material produced (Bankova et al., 1998; Nogueira-neto, 1997). Several compounds such as phenolic acids and water-soluble tannins have been isolated from *Melipona fasciculata* geopropolis (gallotannins and ellagitannins). These compounds, coming from the aqueous and alcoholic fractions of Melipona scutellaris geopropolis (Franchin et al., 2012, 2013), demonstrated antioxidant activity, anti-inflammatory and antinociceptive activities.

Inside the Meliponinae subfamily there is the genus *Scaptotrigona*, which is distributed throughout the Neotropical region and brings together species that build their nests in pre-existing cavities (Silveira et al., 2002). In this genus we have *Scaptotrigona aff postica,* popularly known in Maranhão as the “tubi” bee. However, despite being in the family of geopropolis producers, they are an exception to the rule, as they do not use earth in the composition of propolis (Araújo et al., 2010). Studies on the glandular secretions of bees have demonstrated antibacterial activity, as is the case with the hypopharyngeal secretion of the species Apis mellifera (White, et al., 1963). The same hypopharyngeal glands were found in the species Scaptotrigona postica, assuming that they have the same functions (Costa and Cruz-Landim, 1999).

The vast majority of studies on the pharmacological effects of propolis are related to propolis obtained from *Apis mellifera* bees (Sforcin & Bankova, 2011).There are few studies with *Scaptotrigona aff postica* propolis, and there is no consensus in the literature on how to refer to it, some groups use the name geopropolis (Coelho et al., 2015), while others use propolis (de Farias, et al., 2014; Araújo et al., 2010, 2011). For this work we will use the term propolis, as we believe it to be the most appropriate.

This propolis has been used by the population of the Maranhão region in the treatment of cancer and wound healing (Araújo et al., 2010, 2011). Scientific studies corroborate the possible medicinal effects of this propolis. Martin et al., (2013), treated rats with corneal injuries caused by burns with an emulsion of the crude extract and observed both healing and anti-inflammatory effects. Antitumor activity against Ehrlich’s tumor and reduction in pathology associated with asthma due to an inhibition of migration of inflammatory cells into the alveolar space are also described. In addition to those mentioned, there are no reports on other actions of this product.

Among the best-known activities of propolis, the antiviral action is one of the most notable, as it demonstrates better efficacy against viruses than standard medications (Cornara, et al., 2017). An example of this is the comparative study between an ointment prepared with Canadian propolis, rich in flavonoids, and acyclovir ointment (a drug indicated for the treatment of Herpes simplex, Varicella zoster, Epstein-Barr and Cytomegalovirus viruses). In this study, propolis-based ointment was more effective than acyclovir in healing genital herpetic lesions and reducing local symptoms in the treatment against genital herpes simple x virus (Vynograd, et al., 2000). An in vitro study with Canadian propolis proved to be effective against Herpes simplex, types 1 and 2 during the viral adsorption phase or when incubated directly with the virus. These results indicate that propolis can directly interfere with the virus, preventing the virus from penetrating cells (Bankova et al., 2014). A propolis collected in the city of Moravia, in the Czech Republic, when tested against HVS-1, showed a protective effect when the virus was previously incubated with the extracts (Schnitzler, et al., 2010).

Studies by our group demonstrated the antiviral effect of the crude extract of *Scaptotrigona aff postica* propolis, which led to a 64-fold reduction in the production of picornaviruses and a 32-fold reduction in Influenza viruses (Coelho et al., 2014). The extract also showed antiviral activity against the herpes virus (Coelho et al., 2015). The main constituents found in the total material, flavones, were geopropolis-di-C-glycosides, with a high content of vicenin-2, together with Pyrrolizidine 7 (3-methoxy-2-methyl-butyryl)-9-echimidinylretronecine and caffeoylquinic acid-O-arabinoside (Coelho et al., 2015). However, the studies mentioned above were carried out only with crude hydroalcoho l ic extractions of Scaptotrigona aff postica propolis. The results obtained indicate that propolis and/or its purified components could be a potential source of new antiviral drugs. Therefore, it is important to determine and characterize which components are responsible for the observed activities and test them on other viruses of human interest, especially arboviruses and coronaviruses.

### 1.3 Lonomia obliqua hemolymph

Several studies have demonstrated the presence of bioactive substances in insect hemolymph and their potential use as therapeutic agents. Some of these substances, known as defensins, are part of the insect defense system. Defensins can act as a link between innate and adaptive immunity, and have allowed insects to survive in highly contaminated environments for more than 350 million years, even without them having a developed immune system like vertebrates. However, even without a sophisticated immune system like ours, insects have developed an efficient immune system against parasites and pathogens, which comprises cellular and humoral responses. Cellular mechanisms involve phagocytosis and encapsulation by hemocytes, while humoral responses include the activation of Prophenoloxidase, and the synthesis of antimicrobial peptides by the fat body, which are released into the hemolymph. Two intracellular signaling pathways, Toll and Imd, control the expression of genes encoding antimicrobial peptides. Furthermore, some peptides can act directly on the pathogen, allowing its destruction. During systemic infection, in addition to antimicrobial peptides, other effector molecules are induced and secreted in the hemolymp h, such as stress response proteins, for opsonization, which is the recognition of a foreign body to be phagocytosed, and iron sequestration (Tzou et al., 2002). Our group has been studying the antiviral action present in the hemolymph of *Lonomia obliqua*, a taturana that is quite common in South America and causes many accidents due to contact with its bristles.

In our previous work, the hemolymph of *Lonomia obliqua* was able to reduce the replication of measles, rubeola, poliovurus and influenza viruses (Greco et al, 2009). This hemolymph was fractionated and the fraction with antiviral action was identified. This protein, of approximately 20 kDa, was sequenced and cloned in a baculovirus system as described by Carmo et al, 2012. This recombinant protein reduce herpes virus replication by up to 1,000,000 times (Carmo et al, 2015). This protein was also expressed in a bacterial system, also showing the same antiviral effect when tested against different viruses.

### 1.4 Mygalin

Another substance with great therapeutic potential isolated from arthropods is mygalin, an acylpolyamine, isolated by us from the hemocytes of the spider *Acanthoscurria gomesiana* and identified as bis-acylpolyamine N1, N8-bis (2,5-dihydroxybenzo yl) spermidine (Silva, Daffre and Bulet, 2000, Espinoza, 2020). Mygalin is a molecule with 417 Da, being a potent modulator of innate immune responses (Maffra et al, 2012). In our studies, four molecules with antibacterial properties were isolated, including mygalin (Pereira et al., 2007). This molecule showed antimicrobial action against *E. coli* (Gram-negative), with the minimum inhibitory concentration (MIC) capable of generating 100% growth inhibition of a non-pathogenic *E. coli* strain equal to 85 uM (35.5 ug /mL) from mygalin (Pereira et al., 2007). This substance is probably involved in the immune defense of arachnids, activating iNOS, enhancing nitrite synthesis and increasing the production of TNF-α (Nassar, 2013). As these invertebrates have an exoskeleton (external skeleton) and, in order to grow, they need to leave it, that is, change it from time to time, in a process called ecdysis. At this stage, these arachnids are very exposed to environmental aggressions. Therefore, these antimicrobial molecules, such as mygalin, play an important role at this time.

This substance was synthesized earlier por us (Maffra et al, (2012) and showed a potent antimicrobial activity.

Therefore, like the two previous substances, purified and synthetic mygalin were tested in this work to determine their antiviral action against avian coronaviruses. This study is important not only for a potential product to control this virus in birds of commercial interest, but also to serve as a model for studies against human coronaviruses, since the antiviral mechanisms in different strains of coronavirus are similar.

## 2. **Material and methods**

### 2.1 **Propolis**

The propolis used in this study was obtained from a commercial colony of Scaptotrigona aff Postica, located in the interior of Maranhão, in the Barra do Corda region.

#### 2.1.1. **Aqueous extraction**

The aqueous extraction of propolis was carried out by macerating propolis (45 grams) kept in a freezer at −20 C for 24 hours, until a powder was obtained. This powder was sieved through a stainless granulometric sieve, fine mesh of 0.15 mm, crushed again and sieved again through 1-5 micron sieves. After this procedure, ultrapure water was added to the powder (100mL for every 10 grams) and homogenized under magnetic rotation (IKA® Big Squido - 1,000 rpm) for 24 hours at room temperature protected from light. To separate the precipitate (wax) from the liquid medium (extract), the supernatant was centrifuged at 4°C for 30 minutes at 15,000 rpm (Eppendorf® centrifuge 5804R-). The supernatant was filtered through a 0.22µm millipore membrane, aliquoted and stored in a refrigerator in sterile glass until use.

#### 2.1.2 Propolis Purification

The purification of propolis was performed by chromatography.To This, 1 ml of aquoso extract of propolis foi added to column. The chromatography as performed wth a Phenomenex semi preparative Jupiter 30µM C18 colunm (300A LC column 250×30mm. Absorbance was monitored at 225nm. Flow 8.0 ml/minute was used. All the fractions obtained were tested by their antiviral activity.

### 2.2 Hemolymph of *Lonomia obliqua*

#### 2.2.1 Hemolymph obtention

The hemolymph of *Lonomia obliqua* used in this work was obtained from the hemolymph stock of the Parasitology Laboratory. This hemolymph was originally obtained by the Butantan Institute from caterpillars collected in the municipalities of the extreme west of Paraná, the north of Rio Grande do Sul and the southwest of São Paulo. These caterpillars are part of the animals used to produce antilonomic serum at the Butantan Institute. After removing the bristles with the poison to be used in the production of the antiserum at the Institute, the discarded caterpillars, from the 6th larval instar, had their pseudo-feet cut and the extravasated hemolymph was collected with a Pasteur pipette. After collection, the hemolymph was centrifuged at 1000 g for 10 minutes, inactivated at 56°C for 30 minutes, filtered through a 0.22µm membrane, and stored at –20° C until use.

#### 2.2.2 Synthetic transferin

The transferrin, a protein showing antiviral activity, isolated from *Lonomia obliqua* hemolymph (Carmo et al, 2012), synthesized bu us, was used in this study. The synthetic protein was characterized by mass spectrometry and is identical to native one

### 2.3 Mygalin obtention

Specimens of tarantula spider *A. gomesiana* and hemocytes was acquired as described previously (Silva et al, 2000 and Lorenzini et al 2000). The hemolymph from animals of either sex at differente stages of development was collected from prechille animals by puncture with an apyrogenic syringe. To avoid hemocyte degranulation and coagulation, the hemolymph was collected in the presence of sodium citrate buffer (Silva, Daffre and Bulet, 2000). The hemocytes were remove from plasma by centrifugation ata 80 xg for 10 min at 40C. This procediment was repeat by 2x. The hemocytes collected from hemolymph were homogenized in a Douce appatarus in 2M Acetic Acid. The supernatant, botained by centrifugation ata 1,800 xg fof 30 min at 6oC, was directly subjected to prepurification by solid phase extraction. The Mygalin purified was obtained by RP-HPLC. The purified mygalin and its synthetic chemical analogue were provided by Dr. Pedro Ismael da Silva Junior from the Laboratory for Applied Toxinology at the Butantan Institute. The compound was filtered, aliquoted, protected from light and kept frozen at −80°C until used in the experiments.

#### 2.3.1 Extraction and purification of mygalin from hemocytes

The acidic extract of hemocytes as described earlier by Silva et al, 2000 and Lorenzini et al 2003. To this, the extract were loaded onto three serially linked Sep-Pak C_18_ cartridges (Waters Associates) equilibrated in acidified water (0.046% trifluoroacetic acid (TFA)). The fraction eluted with 40% acetonitrile (ACN) in 0.046% TFA was further concentrated in a vacuum centrifuge (ThermoSavant, 210 model), reconstituted in deionized water (MilliQ, Millipore) and applied to reverse phase high-performance liquid chromatography (RP-HPLC) using a C_18_ semi-preparative column (Vydac™) equilibrated in 2% ACN/0.046% TFA. The bound material was eluted using a linear gradient of 2–60% ACN in 0.046% TFA over 120 min at a flow rate of 1.3 mL/min. The active fraction (AGH1) was further purified to homogeneity by size exclusion chromatography using a Superdex Peptide HR 10/30 (Amersham Biosciences, GE Healthcare). Elution was performed under isocratic conditio ns with 30% ACN in acidified water at a flow rate of 0.5 mL/min. HPLC purification was carried out at room temperature on a Shimadzu LC-10 Ai HPLC system. The column effluent was monitored by ultraviolet (UV) absorbance at 225 nm and the chromatographic fractions were hand-collected, concentrated under vacuum, and reconstituted in deionized water.

#### 2.3.2 Mass spectrometry

MALDI-TOF-MS (matrix-assisted laser desorption/ionization time of flight mass spectrometry) was performed with a Micromass TofSpec S.E. (Waters). Briefly, 1-_μ_L samples in 50% ACN in 0.1% TFA was mixed with 1 _μ_L of saturated matrix _α_-cyano-4-hydroxycinnamic acid solution and approximately 0.4 _μ_L was deposited onto the sample slide and let dry over the bench. Data acquisition and processing were performed by using the MassLynx software. The fragmentation pattern of the mygalin, L-DOPA, and dopamine was investigated by ESI-MS/MS (tandem electrospray ionization mass spectrometry) on a Finnigan LCQ-Duo™ (Thermo Electron Co.). All spectra were acquired in the positive - ion mode, and collision induced dissociation (CID) experiments were performed using a relative collision energy between 20% and 30% (1–1.5 eV).

#### 2.3.3 Synthetic Mygalin

The compound was synthesized by Dr. Pedro Ismael da Silva, Junior from the Laboratory for Applied Toxinology, Butantan Institute as describe by Abraham Espinoza-Culupú et al 2020. The synthetic Mygalin was characterized by tandem mass spectrometry and is identical to native one

### 2.4 **Cell line, culture medium and cell culture**

VERO cells (African green monkey kidney fibroblast cells (Cercophitecus aethiops), were cultured at 37°C in T-flasks or 96-well microplates, containing 5 ml of Leibovitz medium (L-15), supplemented with 10% fetal bovine serum. The cells were maintained at 36°C in a cell culture oven. Cell growth and morphology were monitored daily using an Olympus CK2 inverted optical microscope at 200x magnification. Cell concentration was determined using a hematocytometer and cell viability was determined using the trypan blue cell exclusion method.

A standardized VERO cell bank was produced to carry out the studies. For this purpose, the cells were cultivated in T-75 flasks until confluence was reached. Cells in the exponential phase were resuspended in freezing medium containing 20% Fetal Bovine Serum (FBS), 70% culture medium and 10% sterile glycerin at an approximate concentration of 107 cells/mL. This suspension was aliquoted into cryotubes (Nunc) with a volume of 1.5 mL. The tubes were kept in a −20°C freezer for two hours and then in a −80°C freezer for 24 hours. After this period, the tubes were transferred and kept in liquid nitrogen.

### 2.5 **Cytotoxicity test of samples on cells**

Toxicity testing of all samples was performed on VERO cells. Cells were seeded at a concentration of 104 cells/well in 96-well plates and cultured at 37°C for 24 hours. After 48 hours, the cells were treated with different concentrations of the samples to be tested. PBS was used as a negative control and 10% DMSO as a positive control. After 24, 48 and 72 hours of treatment, cell viability was determined by visualiz ing cell morphology.

### 2.5 Virus and infection

#### 2.5.1 Viruses

The antiviral action of crude propolis from *Scaptotrigona aff postica* and its purified fraction, crude hemolymph from *Lonomia obliqua*, and a recombinant fraction of this hemolymph and mygalin and its synthetic analogue was tested against a strain of avian coronavirus (Beaudette strain). This viral strain is part of the Parasitology laboratory collection and was obtained from the Department. of Preventative Veterinary Medicine, from the Faculty of Veterinary Medicine and Animal Science at USP (kindly provided by Dr.Paulo Brandão).

#### 2.5.2 Viral stock

To carry out this study, a stock batch of the virus was created. To create the viral stock, an aliquot of the viral strain was placed on the cellular mat and left in contact for 1 hour. After this period, the culture medium in the culture bottles was filled with 5 mL of L-15 medium and left in culture for 72 hours to 120 hours, until the onset of the cytopathic effect. After this period, the supernatants were collected and centrifuged to remove residual cells, aliquoted into tubes and frozen at −80°C until the time of the experiments, when they will be titrated to determine the amount of virus to be used in the experiments.

#### 2.5.3 Viral titration

Viral stocks were titrated using the viral extinction technique as described by Reed and Muench, 1938 and detailed in the following item. The viral titer was expressed as TCID50/mL (50% Tissue culture infectious dose), which is the highest dilution of the virus capable of infecting 50% of cells. An aliquot of the virus was thawed immediately before carrying out the experiment.

To determine the viral titer, aliquots of samples of the viruses under study were placed on a mat of VERO cells in semi-confluence (2 x 105 cells/mL), grown in 96 microplates. The viruses were added in serial dilutions ratio 2, As plates were incubated at 36°C and the viral cytopathic effect was observed daily and the titer of infectious viral particles was calculated as described by Reed and Muench, 1938.

#### 2.5.4 Antiviral activity test

##### 2.5.4.1 Antiviral activity test in cell culture (TCID/50)

VERO cells, in exponential growth phase, grown in a 96-well plate, were treated 1 hour before infection or 1 hour after infection, with different amounts of the antiviral agents under tests (2, 5 and 10% for propolis and hemolymph or 26, 52, 104 and 160 µM of mygalin or its analogue). The experiment was also carried out by infecting the cells with a mixture of the virus and the substance under test, kept in contact for 1 hour before being added to the cell culture. The cells were infected with the coronavirus in serial dilutions ratio 2, with the first dilution containing 250 or 1.000 TCID/50 virus. The cultures were maintained at 36°C and observed daily under an optical microscope, seeking to determine the inhibitory effect of the samples on virus replication by determining the cytopathic effect. After the appearance of the cytopathic effect, the culture supernatant was removed and the cells were stained with crystal violet to reveal the foci of cytopathic effect.

## 3. RESULTS

### 3.1. Determination of the cytotoxicity of the agents to be tested

At the beginning of the work, the cytotoxicity of *Scaptotrigona aff. postica, Lonomia obliqua* hemolymph and mygalin were determined in VERO cells. The concentrations used were the same as those used in the tests. For this VERO cells were grown in 96-well microplates. When the cellular confluent was obtained, 1, 2, 5 or 10% v/v of propolis or crude hemolymph, or 26, 52, 104 and 160 µM of mygalin or its synthetic analogue were added to the cultures in triplicate. The crops were observed daily for 72 hours to determine morphological changes in the cell culture. The only component that showed cytotoxic effects in culture was hemolymph at a concentration of 10%, and therefore in the experiments hemolymph was used up to a concentration of 5%. As a negative control for the experiments, in all tests carried out, an aliquot of the agent to be tested was always added to normal VERO cells, at the same concentration as the test and used for comparison with the results obtained.

### 3.2 Determination of the antiviral action of different products on Coronavirus aviary

In this work, the antiviral action of 3 substances (propolis, hemolymph and mygalin and their derivatives) were tested against an avian coronavirus. Despite being harmless to humans, the replication mechanism may be similar to Covid-19, making this virus a model for studying antiviral action, without the need for the entire security structure necessary for Covid-19. For this experiment, 96-well plates were treated with the agents under study, remaining in contact with the cells for 1 hour. After this period, serial dilutions (ratio 2) of avian coronavirus, with an initial titer of 1000 TCID 50, were added to the treated cultures and observed for 72 hours to determine the cytopathic effect.

In **Figure 1** a representative graph of 3 experiments is presented. As can be seen, an intense reduction in the viral effect was observed with all agents tested, mainly with propolis 2% v/v, which had a 16-fold reduction (93.75%).

**Figure 1:**
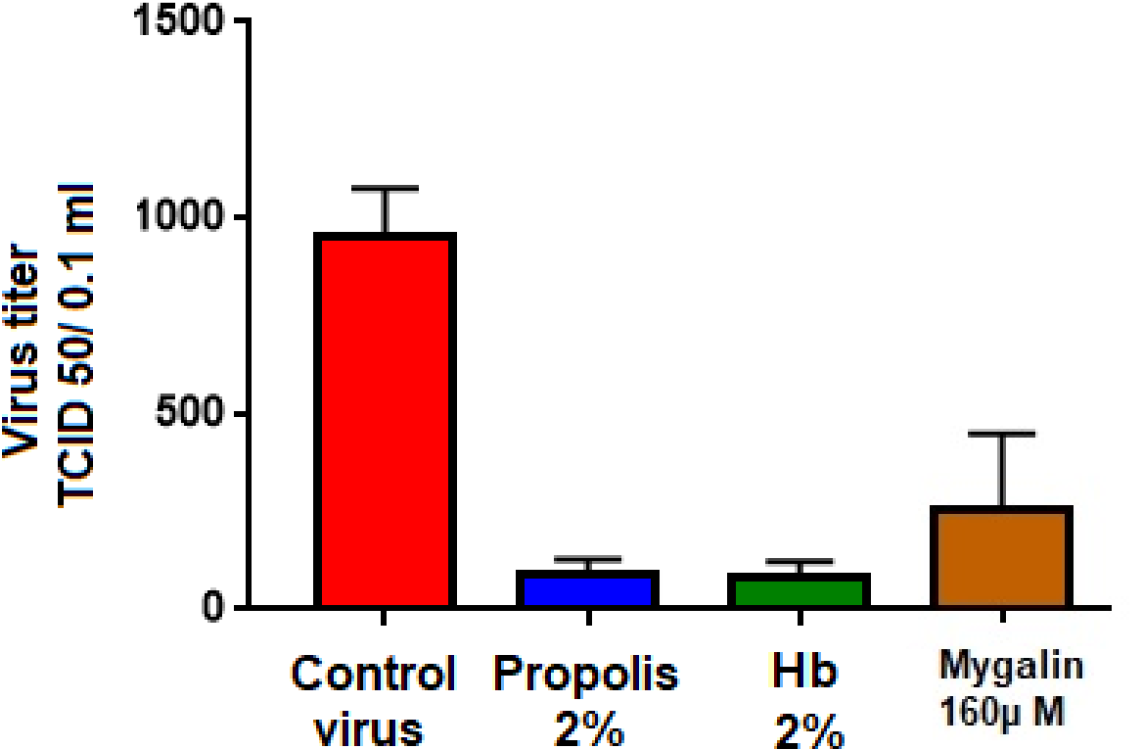
Antiviral action of propolis of *Scaptotrigona aff. postica* (2% v/v) from the hemolymph of *Lonomia obliqua* (2% v/v) or mygalin (160µM). These substances were added to VERO cell cultures 1 hour before the addition of serial dilutions (r:2) of avian coronavirus (initial titer of 1024 TCID 50/0.1ml). The culture was maintained at 36 °C for 72 hours and the culture was observed daily to determine the cytopathic effect. The final titer was determined by the highest dilution of virus causing cytopathic effect in the culture

In **Figure 2**, are presented photos of VERO cells infected with avian coronavirus and treated with different substances under test. As can be seen, there was a clear reduction in the cytopathic effect caused by the avian coronavirus 72 hours after infection, in cultures treated with *Lonomia obliqua* hemolymph (2% v/v) (C), propolis of *Scaptotrigona aff postica* (2% v/v) (D)or mygalin (160 µM) (E). In A is showed a photo of a normal cells culture and in B one culture infected with 250 TCID 50/0.1ml of avian coronavirus.

**Figure 2.**
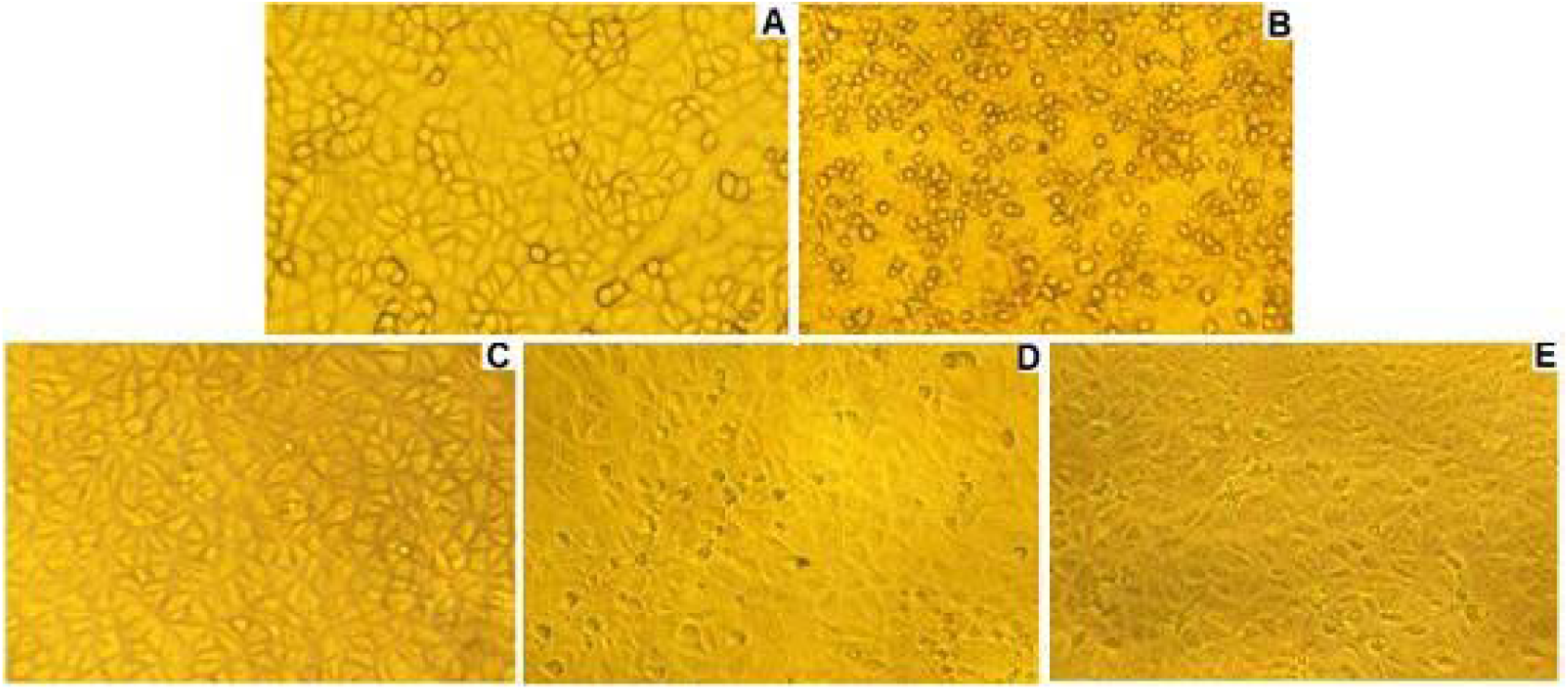
shows a photo of cultures were treated with 2% propolis or 160µM mygalin and infected with 250 TCID 50/0.1ml of avian coronavirus. The cultures were maintained for 3 days at 36°C and after this period the culture medium was removed and the cells were stained with 0.25% crystal violet. The blank excavations were where there was total destruction of the cells by the virus, with no cells attached, while the stained excavations were where the cultures were normal, without cytopathic effects. As can be seen, there was an intense reduction in viral replication both with the addition of propolis and treatment with mygalin.

**Figure 3:**
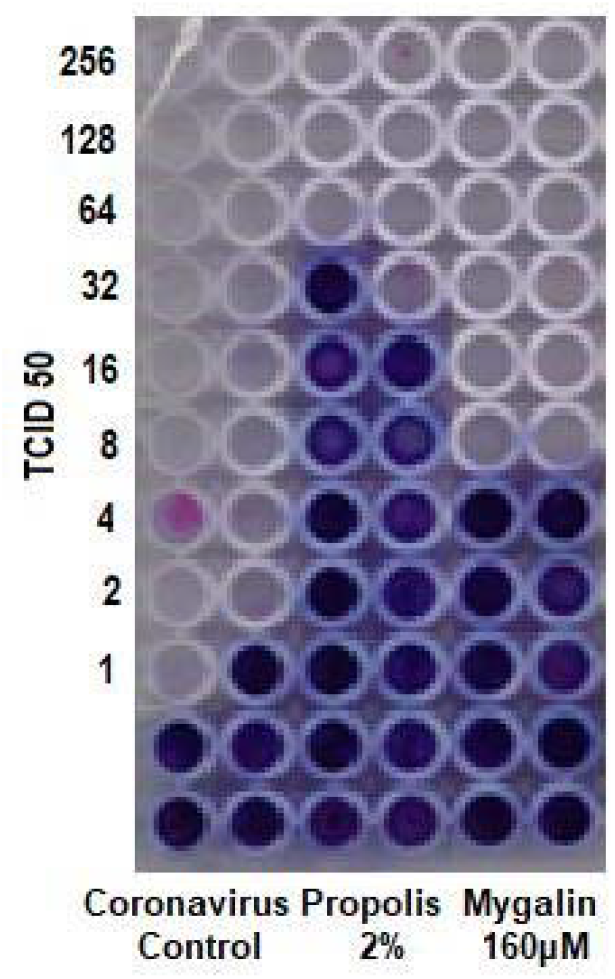
Photomicrograph of a plate of Vero cells infected with 256 TCID 50/0.1ml) of avian coronavirus 1 hour before infection. The culture was treated with 2% propolis or 160µM Migalin. The culture was maintained at 36 °C for 72 hours and the culture was observed daily to determine the cytopathic effect. After that, the cultivation was stained with trypan blue 0.25

#### 3.2.1. Antiviral action of propolis on coronavirus replication

##### 3.2.1.1. Antiviral action with different concentrations of propolis

The antiviral action of different concentrations of propolis (2.5 and 10%) was tested in VERO cell cultures infected with different concentrations of coronavirus (initial titer of 1,000 TCID 50/0.1ml). For this, VERO cell cultures were treated with differe nt concentrations of propolis and after 1 hour these cultures were infected with differe nt concentrations of the virus, in r:2 dilutions. The cultures were maintained at 36 °C for 72 hours and the culture was observed daily to determine the cytopathic effect. The final antivira l activity determined by the protection at the highest dilution of viruses causing a cytopathic effect in the culture. As seen in **figure 4**, the action of propolis is dose dependent, with 2% of propolis being able to reduce the viral titer from 1000 TCID 50/0.1ml to 64 TCID 50/0.1ml, a reduction of 16 times or approximately 94 %. Concentrations of 5 and 10% of propolis were able to reduce virus production to zero in cultures infected with 1000 TCID 50/0.1ml.

**Figure 4:**
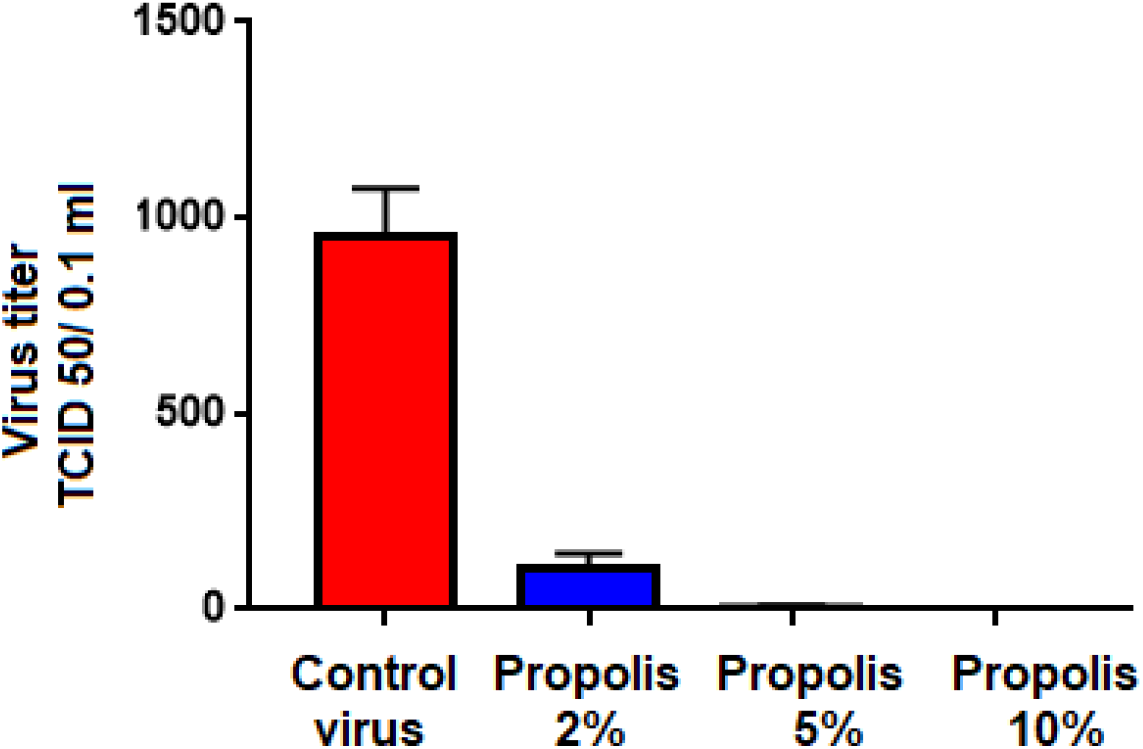
Antiviral action of different concentrations of propolis (2. 5 or 10%) of *Scaptotrigona aff. postica* in the replication of avian coronavirus. These substances were added to VERO cell cultures 1 hour before infection of VERO cells with serial dilutions (r:2) of avian coronavirus (initial titer of 1,000 TCID 50/0.1ml). The culture was maintained at 36 °C for 72 hours and the culture was observed daily to determine the cytopathic effect. The final titer was determined by the highest dilution of virus causing cytopathic effect in the culture

##### 3.2.1.2. Effect of propolis addition time on antiviral action

The effect of the moment of addition of propolis (2%) was tested in VERO cell cultures infected with 1,000 TCID 50/0.1ml of coronavirus. For this, VERO cell cultures were treated with propolis and after 1 hour these cultures were treated 1 hour before infect io n, 1 hour after infection, or with a mixture of viruses plus propolis kept in contact for 1 hour. The cultures were maintained at 36 °C for 72 hours and the culture was observed daily to determine the cytopathic effect. The final antiviral activity determined by the protection at the highest dilution of viruses causing a cytopathic effect in the culture. As seen in **figure 5**, the antiviral effect was more intense when propolis was added 1 hour before infection.

**Figure 5:**
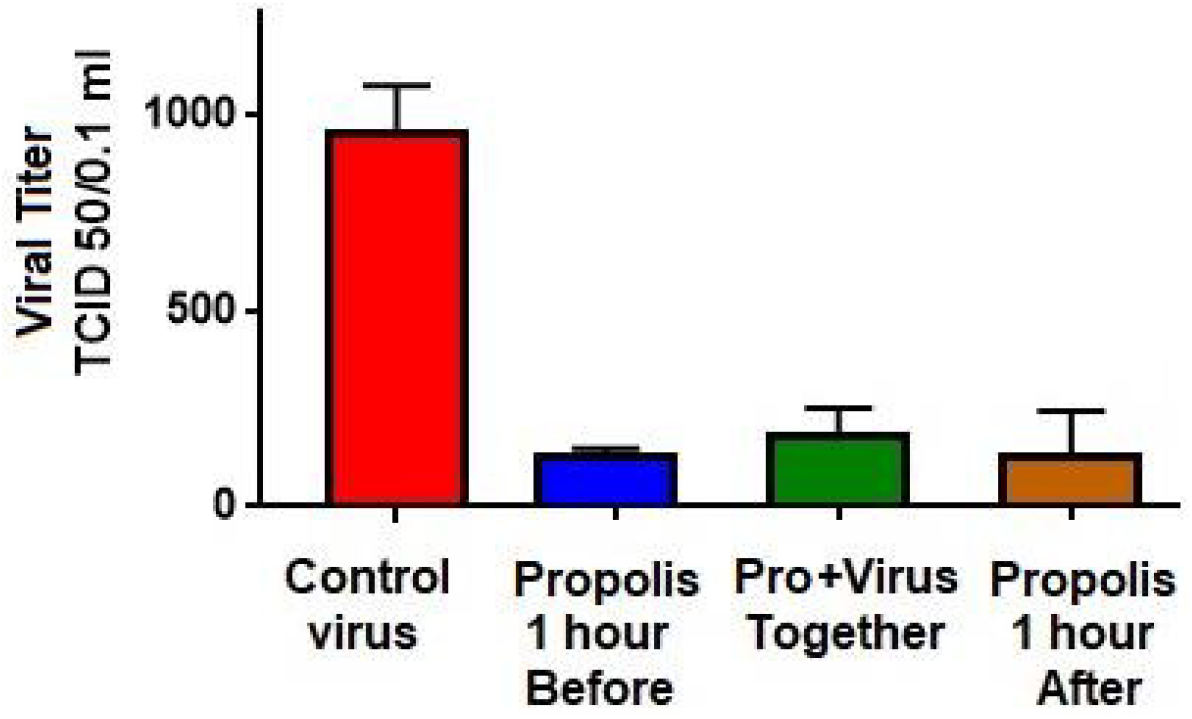
The effect of the time of propolis addition (TOI) was tested in VERO cell cultures infected with 1,000 TCID 50/0.1ml of coronavirus. For this, VERO cell cultures were treated with propolis (2%), 1 hour before infection, 1 hour after infection, or with a mixture of viruses plus propolis kept in contact for 1 hour. The cultures were maintained at 36 °C for 72 hours and the culture was observed daily to determine the cytopathic effect. The final antiviral activity determined by the protection at the highest dilution of viruses causing a cytopathic effect in the culture.

##### 3.2.1.3. Propolis fractionation an antiviral activity determination

To determine which substance present in propolis has an antiviral effect, an aliquot of 1 ml of aqueous propolis was performed by chromatography. To this, 1 ml of aqueous was applied to a Phenomenex semi preparative Jupiter 30µM C18 columnm (300A LC column 250×30mm. Absorbance was monitored at 225nm. Flow 8.0 ml/minute was used. All the fractions obtained were tested for their antiviral activity. As can be seen in the **Figure 6**, the fraction that showed antiviral action was fraction 1, present at the begning of the chromatography. This fraction was capable of reducing viral replication 32 times (from 64 TCID/ 50/0.1ml to 2 TCID/ 50/0.1ml), **Figure 7**.

**Figure 6:**
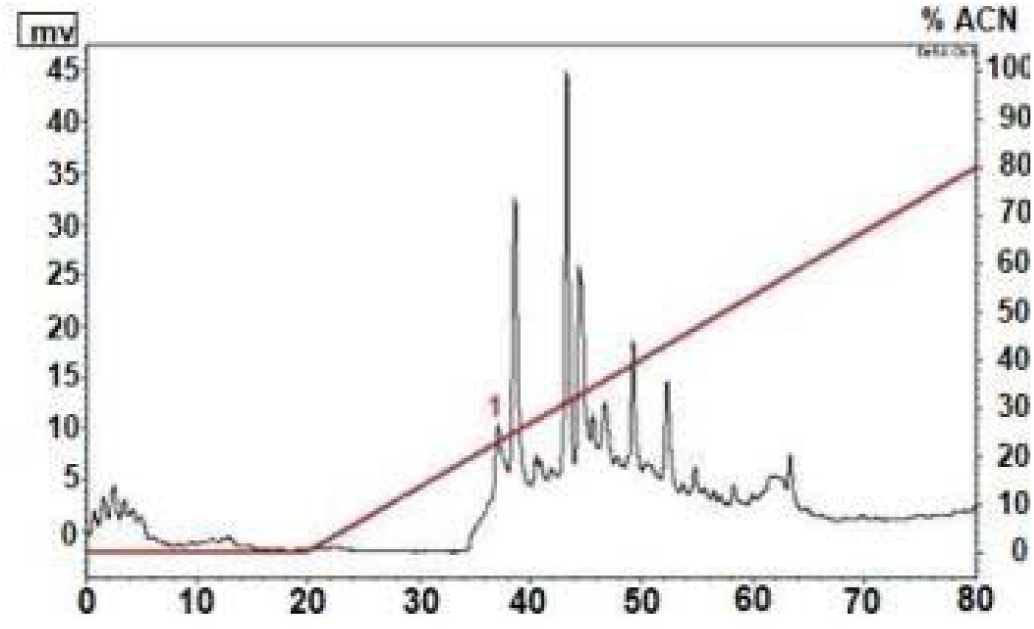
Chromatographic prolife of propolis from *Scaptotrigona aff. postica*. Chromatography was performed with a phenomenex semi preparative Jupiter 30µ C18 columm, 300 LC columm 250×30mm. Absorbance was monitored at 222 nm. Flow of 8 ml/minute was used. All the fractions were tested by their antiviral activity. The fraction 1 as the most active againt avian coronavirus.

**Figure 7:**
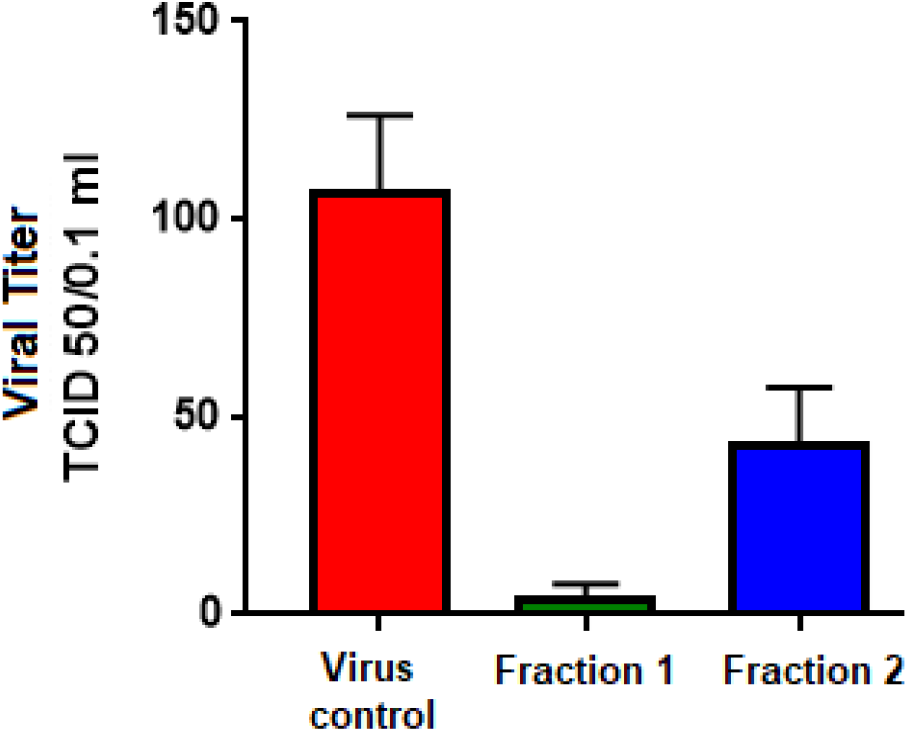
Determination of the fraction of propolis with antiviral action. The antiviral action of the fractions was tested in VERO cell cultures infected with 128 TCID/50/0.1ml of avian coronavirus. For this, VERO cell cultures were treated with propolis fractions (5% v/v), 1 hour before infection. The cultures were maintained at 36 °C for 72 hours and the culture was observed daily to determine the cytopathic effect. The final antiviral activity determined by the protection at the highest dilution of viruses causing a cytopathic effect in the culture.

#### 3.2.2. Antiviral action of hemolymph on coronavirus replication

##### 3.2.2.1 Antiviral action of different concentrations of hemolymph

The antiviral action of different concentrations of *Lonomia obliqua* hemolymph (1.2 and 5%) was tested in VERO cell cultures infected with different concentrations of coronavirus (initial titer of 1,000 TCID 50/0.1ml). For this, VERO cell cultures were treated with different concentrations of hemolymph and after 1 hour these cultures were infected with different concentrations of the virus, in r:2 dilutions. The cultures were maintained at 36 °C for 72 hours and the culture was observed daily to determine the cytopathic effect. The final antiviral activity determined by the protection at the highest dilutio n of viruses causing a cytopathic effect in the culture. As seen in **figure 8**, the action of hemolymph is dose dependent, with 5% of hemolymph being able to reduce the viral titer from 1000 TCID 50/0.1ml to 32 TCID 50/0.1ml, a reduction of 32 times or approximately 97 %. Concentrations of 1 and 2% of hemolymph were also able to significantly reduce virus production in cultures infected with 1024 TCID 50/0.1ml.

**Figure 8:**
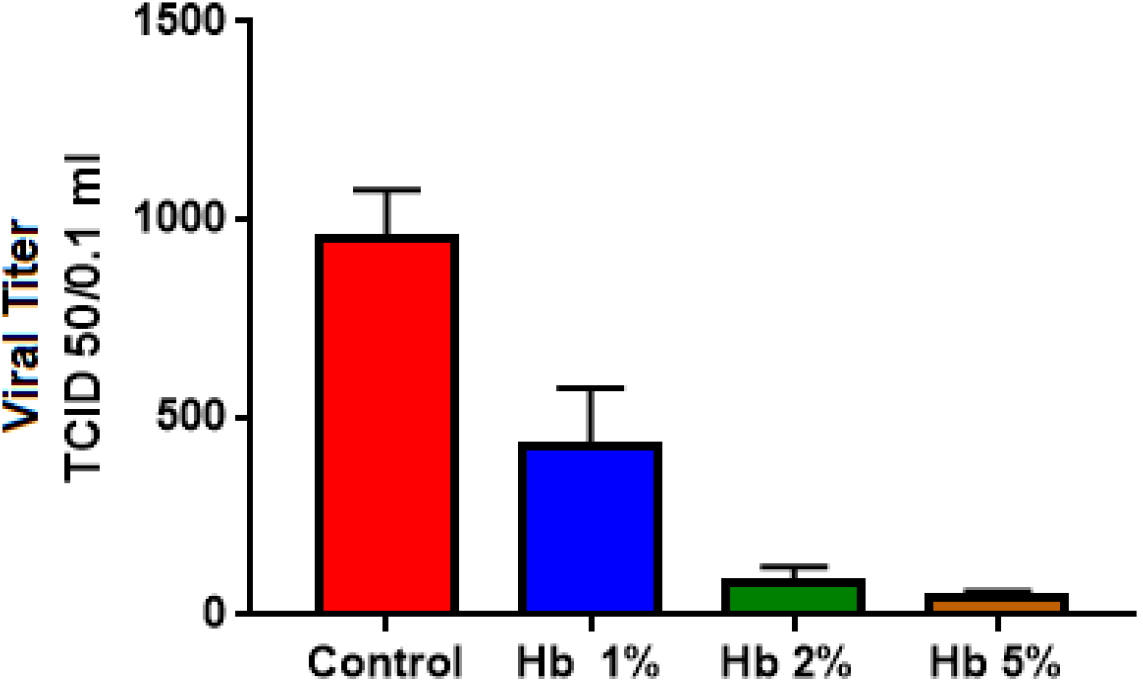
Antiviral action of different concentrations of *Lonomia obliqua* hemolymph (1, 2 and 5%) on the replication of avian coronavirus. Hemolymph were added to VERO cell cultures 1 hour before infection of VERO cells with serial dilutions (r:2) of avian coronavirus (initial titer of 1.000 TCID 50/0.1ml). The culture was maintained at 36 °C for 72 hours and the culture was observed daily to determine the cytopathic effect. The final titer was determined by the highest dilution of virus causing cytopathic effect in the culture.

##### 3.2.2.2 Action of time of addition (TOI) of *Lonomia obliqua* hemolymph on antiviral action

The effect of the moment of addition of *Lonomia obliqua* hemolymph (2%) was tested in VERO cell cultures infected with 1,000TCID 50/0.1ml of coronavirus. For this, VERO cell cultures were treated with hemolymph 1 hour before or infection, 1 hour after infect io n, or treated with a mixture of virus plus hemolymph kept in contact for 1 hour. The cultures were maintained at 36 °C for 72 hours and the culture was observed daily to determine the cytopathic effect. The final antiviral activity determined by the protection at the highest dilution of viruses causing a cytopathic effect in the culture. As seen in **figure 9**, the antivira l effect was more intense when hemolymph was added 1 hour before infection.

**Figure 9:**
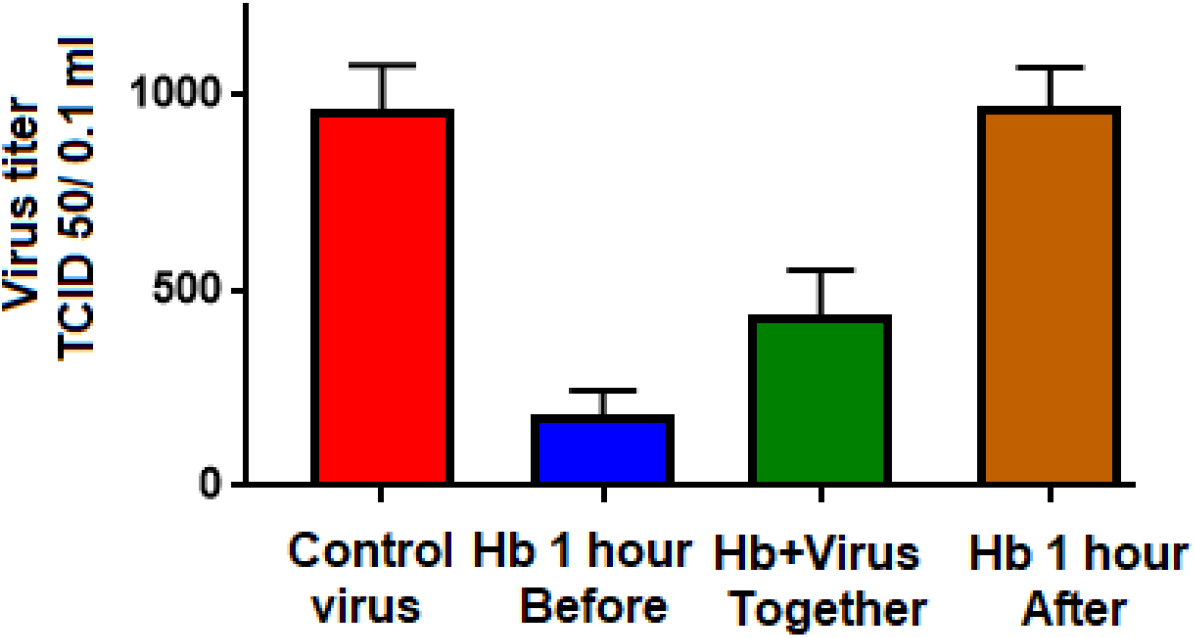
The effect of time of addition (TOI) of *Lonomia obliqua* hemolymph was tested in VERO cell cultures infected with 1,000 TCID 50/0.1ml of coronavirus. For this, VERO cell cultures were treated with hemolymph of *Lonomia obliqua* (2%), 1 hour before infection, 1 hour after infection, or with a mixture of viruses plus propolis kept in contact for 1 hour. The cultures were maintained at 36 °C for 72 hours and the culture was observed daily to determine the cytopathic effect. The final antiviral activity determined by the protection at the highest dilution of viruses causing a cytopathic effect in the culture.

##### 3.2.3.1 Action of adding Transferin on antiviral action

The substance present in the hemolymph, with antiviral action, was identified as a transferin. This substance was synthesized and tested to determine its antiviral action. As can be seen in **figure 10**, synthetic transferin has the same antiviral action observed in crude hemolymph.

**Figure 10:**
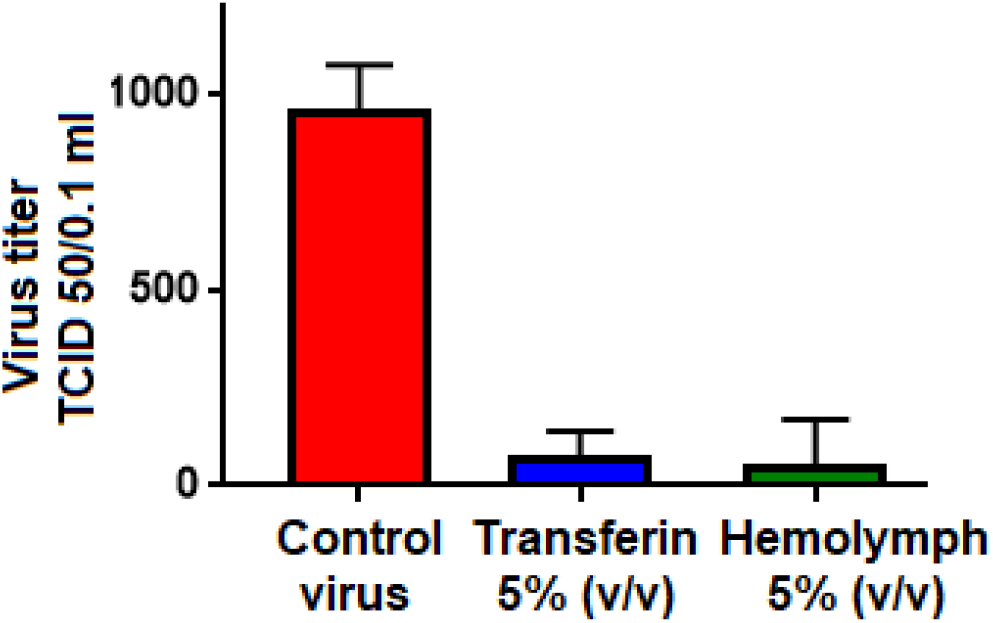
The effect of adding Lonomia obliqua hemolymph, or its synthetic antiviral fraction, was tested in VERO cell cultures infected with 1,000 TCID 50/0.1ml of coronavirus. For this, VERO cell cultures were treated with total hemolymph or its synthetic fraction (tansferin), both 5% v/v, 1 hour before infection. The cultures were maintained at 36 °C for 72 hours and the culture was observed daily to determine the cytopathic effect. The final antiviral activity is determined by protection at the highest dilution of viruses causing a cytopathic effect in the culture.

##### 3.2.3.2 Effect of time of addition of mygalin on antiviral action

The antiviral action of mygalin (160µM) was tested in VERO cell cultures at differe nt times. In the first study, mygalin was added to the culture 1 hour before infecting the cells with serial dilutions (r:2) of avian coronavirus (initial titer of 1.000TCID 50/0.1ml). In the second study, mygalin was kept in contact for 1 hour, with different dilutions of the virus. In the last test, the cultures were infected with different concentrations of virus and after 1 hour mygalin was added. The cultures were maintained at 36 °C for 72 hours and the cultures were observed daily to determine the cytopathic effect. The final antiviral activity determined by the highest dilution of virus causing cytopathic effect in the culture. As seen in **figure 11**, as in previous cases with the other substances tested, the addition of migalin before infect ion was more effective in inhibiting viral replication, but with a less intense effect than with propolis or *Lonomia obliqua* hemolymph.

**Figure 11:**
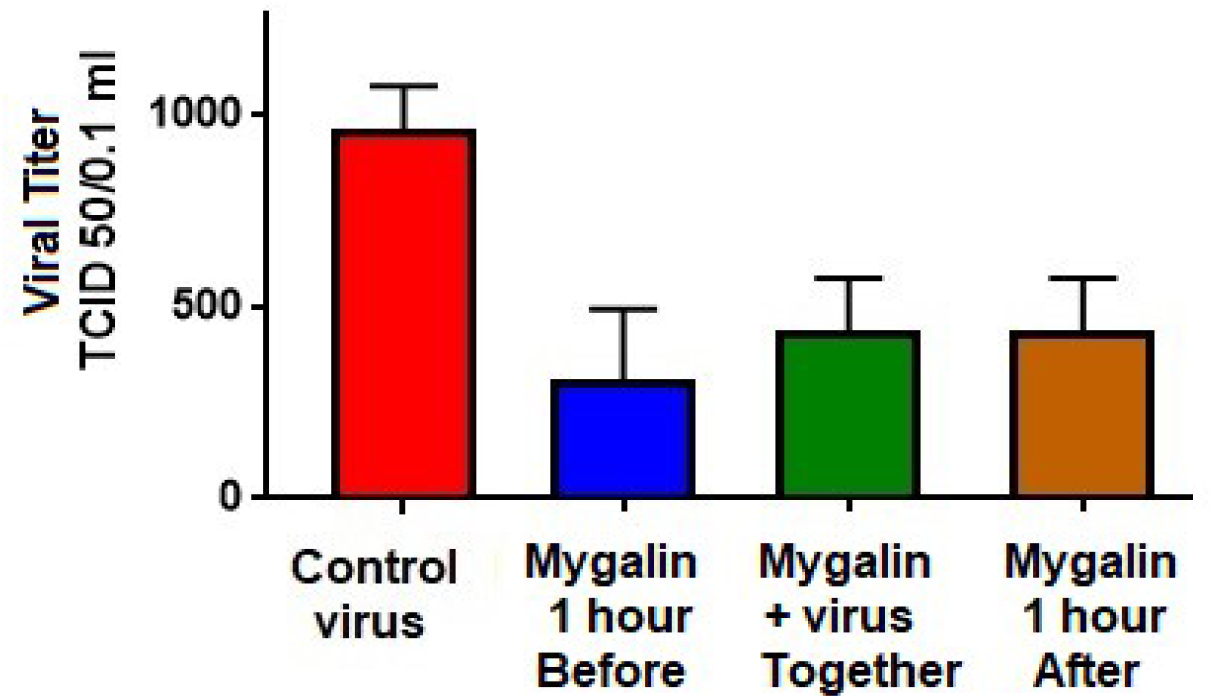
The effect of the time of addition (TOI) of mygalin was tested in VERO cell cultures infected with 1,000 TCID 50/0.1ml of avian coronavirus. For this, VERO cell cultures were treated with 160 µM of mygalin 1 hour before infection, 1 hour after infection, or with a mixture of viruses plus mygalin kept in contact for 1 hour. The cultures were maintained at 36 °C for 72 hours and the culture was observed daily to determine the cytopathic effect. The final antiviral activity determined by the protection at the highest dilution of viruses causing a cytopathic effect in the culture.

##### 3.2.3.3. Effect of synthetic mygalin on antiviral action

The antiviral action of synthetic mygalin was tested. For this, VERO cell cultures were treated with 160 µM of purified mygalin or 26 µM of synthetic mygalin 1 hour before infection with 250 TCID 50/0.1ml of avian coronavirus. The cultures were maintained at 36 °C for 72 hours and the culture was observed daily to determine the cytopathic effect. The final antiviral activity determined by the protection at the highest dilution of viruses causing a cytopathic effect in the culture. The result is shown in **figure 12**. As can be seen, synthetic mygalin is as efficient or more efficient than natural and purified mygalin, extracted from spiders.

**Figure 12:**
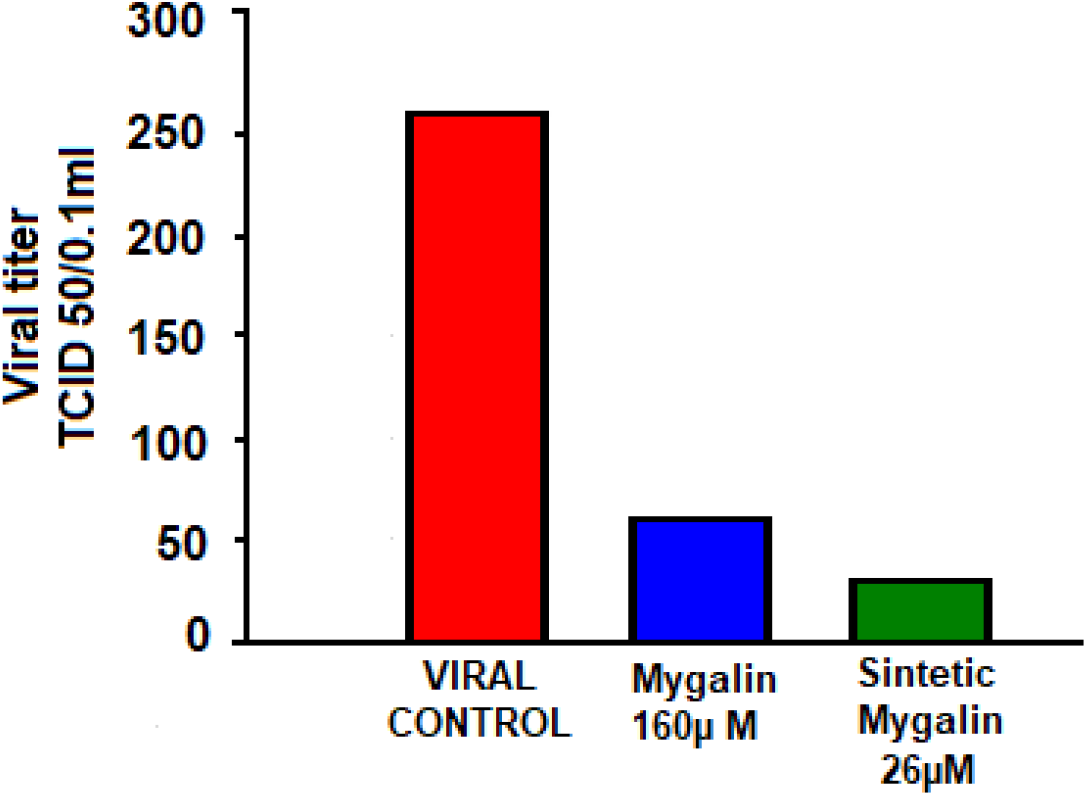
Antiviral action of synthetic mygalin. For this, VERO cell cultures were treated with 160 µM of purified mygalin or 26 µM of synthetic mygalin 1 hour before infection with 250 TCID 50/0.1ml of avian coronavirus. The cultures were maintained at 36 °C for 72 hours and the culture was observed daily to determine the cytopathic effect. The final antiviral activity determined by the protection at the highest dilution of viruses causing a cytopathic effect in the culture.

### 3.3. Antiviral action propolis commercial and from *Scaptotrigona aff* propolis against human coronavirus (Covid-19)

To study whether there is an antiviral action of propolis on human coronavirus (Covid-19) a preliminary experiment was carried out treating VERO cells with propolis of *Scaptotrigona aff. postica* and a propolis of commercial origin (green propolis). After infection, the culture received a layer of agar to prevent the virus from infected cells from spreading throughout the plate. After 72 hours of infection, the agar was removed and the plate stained with 0.25% crystal violet and the presence of viral plaques was observed. The number of viral plaques formed is evidenced by transparent spots on the plaque, caused by the lysis of infected cells. The number of plaques was then counted in control infect ed plaques and in cells infected but treated with propolis, and the result expressed as PFU (plaque forming unit). As can be seen, there is an evident reduction in the number of viral plaques in the experiment carried out with treatment with propolis of *Scaptotrigona aff postica* regarding crops infected with COVID-19, but without treatment **figure 13**.

**Figure 13:**
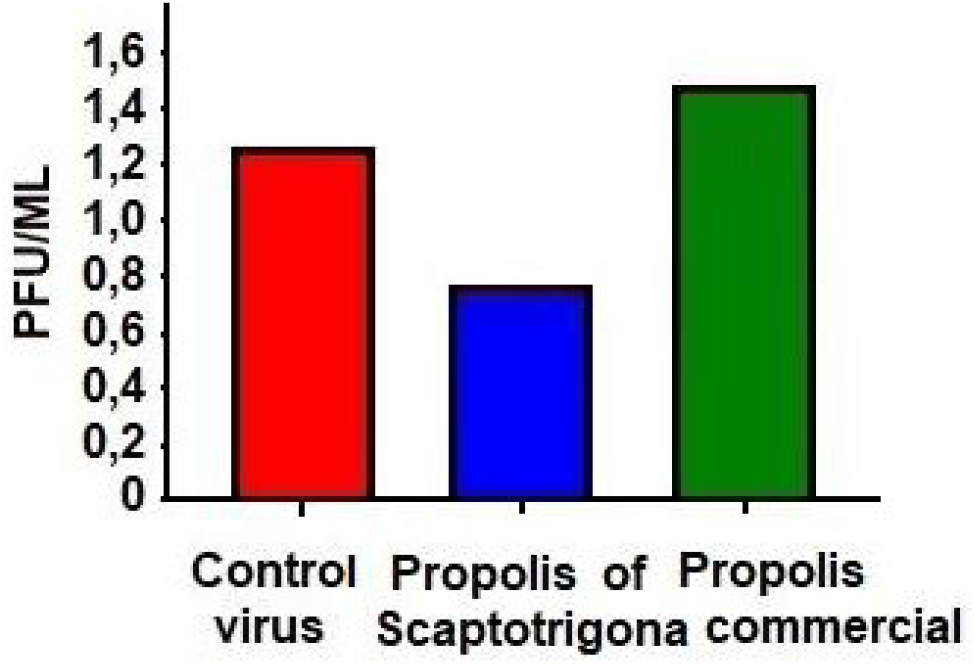
Antiviral action of propolis on human coronavirus (Covid-19). The experiment were carried out treating VERO cells with *Scaptotrigona aff postica* propolis or with propolis of commercial origin (green propolis). One hour after treatment, cells were infected with human Covid-19. After infection, the culture received a layer of agar to prevent the virus from infected cells from spreading throughout the plate. After 72 hours of infection, the agar was removed and the plate stained with 0.25% crystal violet and the presence of viral plaques caused by the lysis of infected cells was observed. The number of plaques was then counted in control infected plaques and in cells infected but treated with propolis, and the result expressed as PFU (plaque forming unit).

## 4. DISCUSSION

Various studies are been performed aiming to identify new substances with potential action antimicrobial in natural source. In this context, arthropods have been an increasingly frequent option. So, the presence of active principles in the arthropods, are of interest for the development of new pharmacological drugs. The antimicrobial protection in arthropods is due to the presence of many peptides, including defensins. Defensins are a major group of antimicrobial peptides and are found widely in vertebrates, invertebrates and plants. In arthropods, these antimicrobial substances can be obtained from differents sources, as hemolymph, muco, saliva, venenos.

Lima-Netto, et al (2012) observed antiviral activity against picornavirus and influe nza virus in the eggs wax from tick *Amblyomma cajennense* (Fabricius). Toledo-Piza et al (2016) have identified an antiviral effect against measles virus in mucus of the land slug *P. Boraceiensis.* Ai et al., (2008, 2012) have found effect on fungus and bacteria on chitosan extracted from M. domestica, while Hou et al., (2007) have observed antibacterial activity in the extract from the larvae of the housefly. Cao X., et al., 2011 have found that this lectin can upregulate NO and iNOS production via TLR4/NF-_κ_B signaling pathway in macrophages. This lectin was was purified by. Earlier we have showed a potent antivira l effect of propolis of *Scaptotrigona aff Postica.* Coelho et al., 2018, have showed action of propolis against rubeola vírus, Carmona et al., 2020 against covid virus and, Mendonça et al, 2022 action antiviral againt arbovirus.

In this present study, all tested substances (propolis, its purified fraction, mygalin, and its synthetic analogue and *Lonomia obliqua hemolymph,* and its synthetic analogue) were capable of inhibiting the avian coronavirus. The results obtained were similar to those obtained in previous experiments for other viruses, suggesting that the mechanism of action may be the same, regardless of the viral type, which can be extremely important as a drug due to its broad spectrum of action.

By the same way, we have shown that hemolymph of *Lonomia oblique* can reduce the viral replication. In earlier study we have observed a reduction of 10^6^ fold in herpes virus replication (Carmo et al., (2012). Despite coronavirus being a virus of different viral family, an intense reduction in virus replication was observed, but not as intensive as to herpesvirus. We agree that this antiviral action of hemolymph can be due to an intracellular interfero n production or activating some other factor of the cells’ innate immune response. This appears to be true because the observed antiviral action of hemolymph is more intense when the substance is isolated from hemolymph is present before infection, probably by stimula t ing the production of some factor linked to the innate antiviral response. For *Lonomia oblique*, this action may also act either on the steps of the cycle of replication of viruses that occur intracellularly, similarly to alloferon, or on the late stages of virus infection, similarly to the peptide from H. virescens (Greco et al., 2009).Furthermore, it seems to be unable either to inhibit the stages before reaching the target cell or to prevent the process of adsorption (Greco et al., 2009). Nevertheles any other antiviral mechanism can be acting in our experiments. As observed with a substance isolated from M. domestica, the antiviral action can be related to some direct deactivating effect against viruses, avoiding the adsorption or penetration of the virus. Therefore, as already described by us, the antiviral action was more intensive when the substance was present earlier virus infection. The protein responsible by antiviral action was determined by us to be a transferin. As can be seen in **figure 10**, synthetic transferin has the same antiviral action observed in crude hemolymph, proving that this substance is responsible for the observed activity.

Another substance, isolated and synthesized by us (Mafra et al., 2012) was the mygalin, an bis-acylpolyamine N1, N8-bis (2,5-dihydroxybenzoyl) spermidine, isolated from the hemocytes of the spider *Acanthoscurria.* This substance, has showed an potent antibacterial action (Pereira et al., 2007) and has demonstrated to be a potent inductor of IFN-gama synthesis and an specific inhibitor of iNOS. Besides thies, mygalin activated macrophages to produce TNF-alfa. So, mygalin can modulate the innate immune response by inducing IFN-gama and NO synthesis. Because these, mygalin was tested by the first time as antiviral effector. As could be observed in this study, mygalin showed a antiviral effect, proving our expectations. Its antiviral action is probably precisely due to the action of these innate immune response effectors described above. The antiviral effect was not as intense as that observed in propolis and *Lonomia obliqua* hemolymph, but this may be related more to the concentration tested than its effector capacity for antiviral action

So, we can say that all the substances tested are a promissor antiviral agents, being of great pharmacological interest, since that, an intense reduction in viral replication was obserded, at the doses used here, and without presenting a toxic action on cells.

## Acknowledgments

We thank everyone involved in the work.

## Ethical Approval

This work only involves invertebrate organisms and has ethical approval for its development. We emphasize that the work does not involve vertebrate animals.

## Consent to Participate/Consent to Publish

All authors have read and agreed to the published version of the manuscript.

## Authors Contributions

All authors had equal participation in the development of the work.

## Availability of data and materials

The authors declare that data and materials are available upon request.

